# Unveiling the Mechanisms to Bypass KRAS Inhibition: *In Vitro* Insights into the Influence of Fibroblast-Secretome

**DOI:** 10.1101/2024.04.19.590324

**Authors:** Susana Mendonça Oliveira, Patrícia Dias Carvalho, André Roma, Patrícia Oliveira, Andreia Ribeiro, Joana Carvalho, Flávia Martins, Ana Luísa Machado, Maria José Oliveira, Sérgia Velho

**Author notes:** Corresponding author: Sérgia Velho.

## Abstract

Novel KRAS-targeted therapies unlocked new treatment options for previously untreatable patients. However, in colorectal cancer (CRC), resistance to KRAS-targeted therapy develops rapidly, making it imperative to understand its underlying mechanisms.

Cancer-associated fibroblasts (CAFs) induce therapy resistance by generating and maintaining cancer stem cells (CSCs). Additionally, CAFs secretome can modulate KRAS mutant CRC cells proteomic profile, independently of mutant KRAS. Hence, we investigated whether CAF-derived factors could induce resistance to KRAS inhibition by promoting a KRAS-independent stem-like phenotype.

Evaluation of KRAS-mutant CRC cell lines (HCT15, HCT116, and SW480) revealed unique basal stem cell marker expression levels. Silencing KRAS lead to up-regulation of CD24, down- regulation of CD49f and CD104, and reduced stemness. However, CAF-secreted factors attenuated these effects, restoring stem cell markers expression and increasing stemness. RNA sequencing showed that CAF-secreted factors upregulate pro-tumorigenic pathways in KRAS-silenced cells, including cell cycle control, epithelial-mesenchymal transition (EMT), NOTCH, and immune regulation, leading to increased cell cycling and exit from quiescence.

Overall, we provide mechanistic insights illuminating the role of fibroblasts in counteracting KRAS silencing-induced growth inhibition and enhancing stemness. Our results show that the limited success of KRAS-targeted therapies is not only derived from cell-intrinsic factors but also dependent on external factors derived from the tumor microenvironment, thus opening avenues to improve therapy responses in CRC.

## Introduction

For over three decades, KRAS oncogene has gained a reputation of an undruggable target (Patricelli et al., 2016; Salgia et al., 2021). However, recent breakthroughs in developing allele-specific covalent inhibitors have paved the way for a new era of KRAS-targeted therapies (Moore et al., 2020; Patricelli et al., 2016). Novel KRAS-targeted therapies are currently in clinical trials, showing promising results, although with a lower response rate in CRC (Huang et al., 2021). Still, patients quickly start to develop drug resistance mechanisms, limiting the effectiveness of the treatment (Akhave et al., 2021).

Among the various factors contributing to the acquisition of resistance to the therapy, the tumor microenvironment plays a pivotal role, in part, due to the presence of CSCs and a supportive network of CAFs (Hu et al., 2015; Prieto-Vila et al., 2017; T. Wu & Dai, 2017).

CSCs harbor both self-renewal potential and the capacity to originate high proliferative, short-lived heterogeneous progeny that compose the tumor bulk. These characteristics, coupled with their intrinsic resistance to treatments, make them accountable for tumor recurrence (Clara et al., 2020).

This is also the case in CRC, where RAS isoforms selectively control the expansion of CSCs, with the KRAS isoform having more potential to promote stemness characteristics in cells. KRAS activation can induce stemness potential through the upregulation of pathways such as the Wnt/β-catenin and the Hedgehog pathways and the increase of surface markers of stemness (Chippalkatti & Abankwa, 2021).

In CRC, cancer stem cells can be distinguished and isolated from tumors based on the expression of surface markers such as CD133, CD44 and its isoform CD44v6, CD166, CD24, CD49f, EpCAM and LGR5 (Barker et al., 2007; Dalerba et al., 2007; Haraguchi et al., 2013; O’Brien et al., 2007; Ricci-Vitiani et al., 2007; Todaro et al., 2014). Still, there is no consensus on the biomarker or biomarker combination that better defines colorectal CSCs. Whether in some works CD133 seems to be enough to identify CRC cells with stem-like properties (Ricci-Vitiani et al., 2007), others suggest EpCAMhigh/CD44+ (Dalerba et al., 2007), CD49f+ (Haraguchi et al., 2013) or Lgr5+CD44+EpCAM+ (Leng et al., 2018) as more reliable indicators of stem potential. Nonetheless, all studies show that CRC possesses a rare cell population that resembles CSCs, regardless of the markers used for isolation, and that they preferentially localize in areas enriched in CAFs (Lenos et al., 2018). In fact, consensus molecular subtype 4, one of the four molecular subtypes identified in CRC, is enriched in CAFS and shows upregulation of genes related to cancer stem cells, as well as EMT, angiogenesis, the complement-mediated inflammatory system, angiogenesis and transforming growth factor beta (TGF-β) pathways (Guinney et al., 2015).

Interestingly, it was also shown that CSCs, as a subpopulation of cells within the tumor tissue, might be precursors to CAFs (Nair et al., 2017). On the other hand, CAFs are responsible for the induction and maintenance of cancer stem cell phenotype through their secreted factors (Linares et al., 2021; Valcz et al., 2018; F. Wu et al., 2021; Yu et al., 2014), highlighting the co-dependence of both these cell types.

Besides regulating the cancer stem cell phenotype, CAFs can also alter the proteome profile associated with CRC cells harboring a KRAS mutation. Specifically, we showed that KRAS oncogenic signaling is predominantly governed by fibroblasts, though most of the fibroblast-associated signaling is independent of mutant KRAS (Dias Carvalho, Martins, et al., 2022).

Herein, we hypothesized that CAF-derived factors might be responsible for driving the stem-like phenotype either dependent or independent of mutant KRAS, subsequently inducing resistance to KRAS-targeted inhibition. Our results confirmed that both KRAS and recombinant human (rh) TGFβ1-activated fibroblasts-secreted factors enhance CRC stem cell activity. Furthermore, they also revealed that rhTGFβ1-activated fibroblasts-derived factors recover the stemness potential lost upon KRAS silencing and lead to a more mesenchymal phenotype through the upregulation of EMT and pro-tumorigenic pathways. These results are extremely relevant as they highlight a potential mechanism of resistance to KRAS-target therapies, independent of KRAS and mediated by the secretome of fibroblasts.

## Materials and methods

### Cell culture

Human CRC cell lines HCT-116, HCT-15, SW480 (table 1), and normal human intestinal fibroblasts CCD18-Co cell line were purchased from the American Type Culture Collection (ATCC). Cells were routinely maintained at 37°C in a humidified atmosphere with 5% CO_2_ in the recommended media: RPMI-1640 media (Gibco, Thermo Fisher Scientific, USA) for the CRC cell lines and DMEM (Gibco, Thermo Fisher Scientific, USA) for the fibroblasts, both supplemented with 10% heat-inactivated fetal bovine serum - FBS (Hyclone, USA) and 1% penicillin-streptomycin - P/S (10,000 U/mL; Gibco, Thermo Fisher Scientific, USA).

**Table 1.** Genetic and histological characterization of KRAS mutant CRC cell lines.

| Cell line | KRAS mutation | Disease | Derived from |
| --- | --- | --- | --- |
| HCT116 | G13D | Colorectal Carcinoma | Primary tumor |
| HCT15 | G13D | Colorectal Adenocarcinoma | DLD-1 misclassified |
| SW480 | G12V | Colorectal Adenocarcinoma | Primary tumor |

### Production of conditioned media of fibroblasts

Fibroblasts were plated into T75 culture flasks and grown in DMEM supplemented with 10% heat-inactivated FBS and 1% P/S and allowed to grow at 37°C in a humidified atmosphere with 5% CO_2_ until approximately 90% of confluence. After washing two times with Phosphate-buffered saline (PBS), the media of the fibroblasts was changed to DMEM supplemented with 1% P/S plus 10ng/mL rhTGF-β1 (ImmunoTools, Germany). The addition of rhTGF-β1 in the media leads to the activation of the fibroblasts, conferring them a CAF-like phenotype.

For the sphere formation experiments, the fibroblasts conditioned media was prepared with DMEM without phenol red (Gibco, Thermo Fisher Scientific, EUA) in the same conditions.

After 4 days in culture, the conditioned media was harvested, centrifuged at 1200 revolutions per minute (rpm) for 5 minutes, filtered through a 0.2μm filter, and stored at -20°C. The cells were harvested with 0.05% Trypsin-EDTA (Gibco, Thermo Fisher Scientific, USA), counted and total protein extraction was performed. The confirmation of fibroblasts activation was assessed through the evaluation of alpha-smooth muscle actin (α-SMA) expression by western blotting.

### Cell culture with conditioned media

Cells were plated in a 6-well plate in a confluence of 150 000 cells per well. After 16 hours the cells were transfected. After 6 hours of transfection, the conditioned media of activated fibroblasts (grown in DMEM + 1% P/S + 10ng/mL rhTGF-β1) was added. After 48 hours in the conditioned media (total of 72 hours of transfection) the cells were harvested with 0.05% Trypsin-EDTA (Gibco, Thermo Fisher Scientific, USA), counted, and collected for cytometry and total protein extraction. KRAS silencing efficiency was assessed by western blotting.

### siRNA Transfection

Knockdown of KRAS was achieved by gene silencing using a pool of 4 small interfering RNAs (siRNA) specific for KRAS (ON-TARGETplus SMARTpool, from Dharmacon, GE Healthcare, USA). The desired cell line was plated onto a 6-well plate and allowed to adhere (150 000 cells per well). The following day, transfection was carried out. The siRNA mix was prepared as following: 250μL of opti-MEM medium (Gibco, Thermo Fisher Scientific, USA) and 3μL of Lipofectamine RNAiMAX (Invitrogen, Thermo Fisher Scientific, USA) were gently mixed and incubated for 5 minutes at room temperature. A mixture of 1μL of siRNA and 250μL of opti-MEM was added to the previous solution and incubated for an additional 20 minutes. During that time, the culture media was completely removed and 1.5 mL of opti-MEM was added to each well. After the incubation period, 500μL of the siRNA mix were added per well. siRNA final concentration was 10nM. As a control, a condition using non-targeting siRNA (ON-TARGETplus Non-targeting Control siRNA #1, from Dharmacon, GE Healthcare, USA) was used at the same concentration as the siRNA targeting the genes of interest. KRAS silencing efficiency was assessed by western blotting. Total silencing time was 72 hours.

### Protein extraction and western blotting

Cells were lysed using RIPA lysis buffer [50mM TrisHCl pH 7.5, 1% (v/v) IGEPAL CA-630, 150mM NaCl and 2mM EDTA] supplemented with 1:7 protease inhibitor cocktail (Roche Diagnostics GmbH, Switzerland) and 1:100 phosphatase inhibitor cocktail (Sigma Aldrich, USA). Cells were centrifuged at 14000 rpm at 4°C, during 10 minutes. The supernatants were collected and stored at -20°C. Protein concentration was determined using the Bradford assay (Bio-Rad Protein Assay kit, Bio-Rad, USA). Equal amounts of protein from each sample were dissolved in sample buffer [Laemmli with 5% (v/v) 2-β-mercaptoethanol and 5% (v/v) bromophenol blue] and denaturated for 5 minutes at 95°C. Samples were separated by 12% sodium dodecyl sulfate-polyacrylamide gel electrophoresis (SDS– PAGE) and proteins were transferred into nitrocellulose membranes (Amersham Protran Premium 0.45μm nitrocellulose blotting membranes, GE Healthcare Life Sciences, USA). For immunostaining, membranes were blocked with 5% (w/v) non-fat dry milk in PBS containing 0.5% (v/v) Tween20 (Sigma Aldrich, USA), and primary antibodies were incubated overnight at 4°C with agitation (table 2). After washing five times with PBS-Tween20 for 5 minutes, membranes were incubated with HRP-conjugated anti-mouse secondary antibodies (Ge Healthcare, USA) for 1 hour at room temperature. Membranes were washed again five times with PBS-Tween20 for 5 minutes. Proteins were developed using ECL blotting substrate (Clarity Western ECL Substrate, Bio-Rad, USA). ImageJ software was used for protein quantification.

**Table 2.** List of antibodies used for western blotting.

| Antibody | MW (kDa) | Blocking | Dilution | Species | Brand | Reference | Storage (°C) | Secondary Dilution |
| --- | --- | --- | --- | --- | --- | --- | --- | --- |
| α-SMA | 42 | 5% Milk in PBS-0.5%T | 1:250 | Mouse | Abcam | ab7817 | -20 | 1:3000 |
| GAPDH | 37 | 5% Milk in PBS-0.5%T | 1:10000 | Mouse | SantaCruz | sc-47724 | 4 | 1:20 000 |
| KRAS | 21 | 5% Milk in PBS-0.5%T | 1:4000 | Mouse | LSBio | C175665 | -20 | 1:8000 |
| HSC70 | 70 | 5% Milk in PBS-0.5%T | 1:8000 | Mouse | SantaCruz | sc-7298 | 4 | 1:16 000 |

MW - molecular weight | kDa - kilodalton

### Flow cytometry

For flow cytometry analysis, cells were harvested using trypsin and resuspended in RPMI supplemented with 10% heat-inactivated FBS and 1% P/S to inactivate the trypsin (completed media). Cells were allowed to recover their membrane markers by a 20-minute incubation in completed media, in an incubator at 37°C and a humidified atmosphere with 5% CO_2_. For the labeling, 200 000 cells were used per condition. Cells were washed with wash buffer (0.5% FBS in PBS) and resuspended in that same wash buffer. Single-cell suspension was labeled using a 1:100 concentration (v:v) of antibody in wash buffer (table 3). As a control, a condition where no antibodies were added to the wash buffer was used (unstained). Fluorochrome-conjugated antibodies were incubated at room temperature, in the dark, for 15 minutes. Labeled cells were then rinsed in wash buffer and finally resuspended in PBS. Cells were analyzed using a FACS Canto-II or BD Accuri C6 (BD Biosciences) flow cytometer. The gating strategy used can be found as supplementary data (Supplementary Figure 1). Data was examined using FlowJo cytometry analysis program. After performing a doublet exclusion gate, both the percentage of positive cells and the median intensity fluorescence were analyzed.

**Table 3.** List of anti-human antibodies used for flow cytometry.

| Antibody | Conjugate | Clone | Reference | Brand |
| --- | --- | --- | --- | --- |
| CD24 | PE | 32D12 | 130-098-861 | Miltenyi Biotec |
| CD44 | FITC | DB105 | 130-113-896 | Miltenyi Biotec |
| CD44.V6 | APC | REA706 | 130-111-425 | Miltenyi Biotec |
| CD49f | APC | GoH3 | 130-100-147 | Miltenyi Biotec |
| CD104 | FITC | REA236 | 130-124-266 | Miltenyi Biotec |
| CD133 | PE | AC133 | 130-098-826 | Miltenyi Biotec |
| CD166 | APC | REA442 | 130-106-619 | Miltenyi Biotec |

### Sphere forming assay

Following siRNA transfection, cells were enzymatically harvested using Trypsin and resuspended in RPMI supplemented with 10% heat inactivated FBS and 1% P/S. Cells were centrifuged at 1200rpm for 5 minutes, the supernatant was removed, and the pellet was washed with PBS. Cells were again centrifuged at similar conditions and resuspended in DMEM without phenol red and 1% P/S. Single-cell suspension was achieved by physical dissociation with a 25-gauge needle. Cells were then plated at a density of 500 cells/cm2 into 6-well plates coated with 1.2% poly(2-hydroxyethylmethacrylate) (Merck, Germany) in 95% ethanol (Sigma-Aldrich, USA) to allow non-adherent culture conditions. Cells were allowed to grow for 5 days in DMEM without phenol red containing 1x B27 supplement (Life Technologies, USA), 1x N2 supplement (Life Technologies, USA), 20 ng/ml EGF (Sigma-Aldrich, USA), 10 ng/mL bFGF (Life Technologies, USA), and 1% P/S in an incubator at 37°C in a humidified atmosphere with 5% CO_2_. For the conditions where fibroblast conditioned media was used, it was only added 1x B27 and 1x N2 supplement because fibroblasts produce growth factors on their own (Kurobe et al., 1985; Powers et al., 2000; Pratsinis & Kletsas, 2016). Sphere forming efficiency (SFE) was calculated as the number of spheres (≥ 50 μm) formed divided by the number of cells plated and multiplied by 100 to be expressed as a percentage 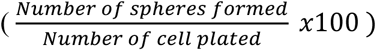. To observe and acquire pictures of the spheres (in brightfield), the IN Cell Analyzer 2000 (GE Healthcare, EUA) microscope was used. To automatically identify the spheres in the images, ilastik: interactive machine learning for (bio)image analysis was employed, as well as the Cell ProfilerTM cell image analysis software. Finally, each dataset was manually curated using the ImageJ software.

### RNA extraction

RNA was extracted from the cells lysed with RLT+ buffer using the RNeasy Mini kit (Qiagen, Germany) according to the manufacturer’s instructions. RNA volume and purity were measured using the UV-Vis spectrophotometer NanoDrop 1000 (Thermo Fisher Scientific, EUA). RNA was stored at -80°C until required.

### Library preparation and transcriptome sequencing

Spheres of KRAS silenced cells formed in the conditioned media of activated fibroblasts were compared with their KRAS silenced counterpart formed in control media. Due to the low sphere-forming efficiency, the quantity of RNA extracted from each experiment was low, especially for the control condition. Therefore, a pool of 3 biological replicates was used per condition. The samples were processed using Ion Torrent technology. Libraries were prepared for each sample with the Ion AmpliSeq Transcriptome Human Gene Expression Kit targeting 20802 genes and sequenced in a 540-chip using the Ion 540 Kit-Chef and the S5 XL instrument (IonS5XL, Thermo Fisher, EUA). The average mean read length was 113 base pairs. The produced reads were aligned to hg19 AmpliSeq Transcriptome ERCC v1 reference sequence and hg19_AmpliSeq_Transcriptome_21K_v1 target regions.

### Database for Annotation, Visualization and Integrated Discovery (DAVID)

Differential expression was analyzed with the help of DAVID online software (https://david.ncifcrf.gov/home.jsp). This comprehensive approach allowed us to perform a preliminary gene annotation and visualization to explore the biological functions and signaling pathways associated with both our gene sets and create tables of functional gene enrichment.

### Gene set enrichment analysis (GSEA)

Gene expression data was analyzed for enrichment using GSEA software (Broad Institute-version 4.1.0) and Human MSigDB v2023.1.Hs following the guidelines of Liberzon et al., 2015; Mootha et al., 2003; Subramanian et al., 2005. Several human collections from the Molecular Signatures Database were used as the gene sets of interest, enabling us to explore the enrichment of biologically relevant pathways and functional annotations. The normalized enrichment score (NES) for each gene was calculated and used for further analysis and graphical representation.

### Quantification of G0 arrest

The quantification of the G0 arrest in the RNAseq data was adapted from the G0 arrest score quantification described in Wiecek et al., 2023. R software was used for gene enrichment and determination of G0 arrest score (original code available in: https://github.com/secrierlab/CancerG0Arrest/tree/main/TCGA_QuiescenceEvaluation). A final positive G0 arrest score indicates that cells are quiescent, while if negative, cells are cycling.

### Statistical Analysis

Results are representative of three or more independent experiments. Quantifications are expressed as mean ± standard deviation (SD) of the biological replicates considered. Statistical analyses were performed using GraphPad Prism v8 (GraphPad Software Inc., USA). All samples were tested for normality and significant statistical difference was considered when the *P* value was less than 0.05. The symbols *, **, *** and **** were used to denote levels 0.05, 0.01, 0.001 and 0.0001 of statistical significance, respectively. The statistical tests performed for each analysis are indicated in the corresponding figure legend.

## Results

### CRC cells express variable basal levels of cancer stem cell markers

In order to ascertain the effects of mutant KRAS in modulating the cancer stem cell landscape in the context of CRC, we selected three CRC cell lines (HCT116, HCT15, and SW480), all harboring a mutation for KRAS (table 1) but with different origins and genetic profiles. As an initial study to characterize the basal stemness potential of each cell line, flow cytometry was performed for the identification of the most commonly used stem cell markers: CD24, CD44 and its isoform CD44v6, CD133, and CD166 (Table 3). Furthermore, we also analyzed integrins that are commonly deregulated upon KRAS oncogenic activation, namely CD104 (Integrin β4) and its binding partner CD49f (Integrin α6) (S. H. Choi et al., 2022; Haraguchi et al., 2013; Zhang et al., 2017). CD49f has been described as a biomarker transversally present amongst stem cell populations, including the intestinal one (Krebsbach & Villa-Diaz, 2017), and as a possible regulator of stem cell markers, such as CD44 (Desgrosellier & Cheresh, 2010). Therefore, CD49f and CD104 were hereafter included in the general panel of cancer stem cell markers.

Our results show that the basal expression of cancer stem markers in the membrane of CRC cell lines is heterogeneous, both in the percentage of positive cells and in the level of expression per cell, denoted by the median fluorescence intensity (MFI), within and across cell lines (figure 1). Only CD49f and CD104 were highly and consistently expressed across all the CRC cell lines.

**Figure 1:**
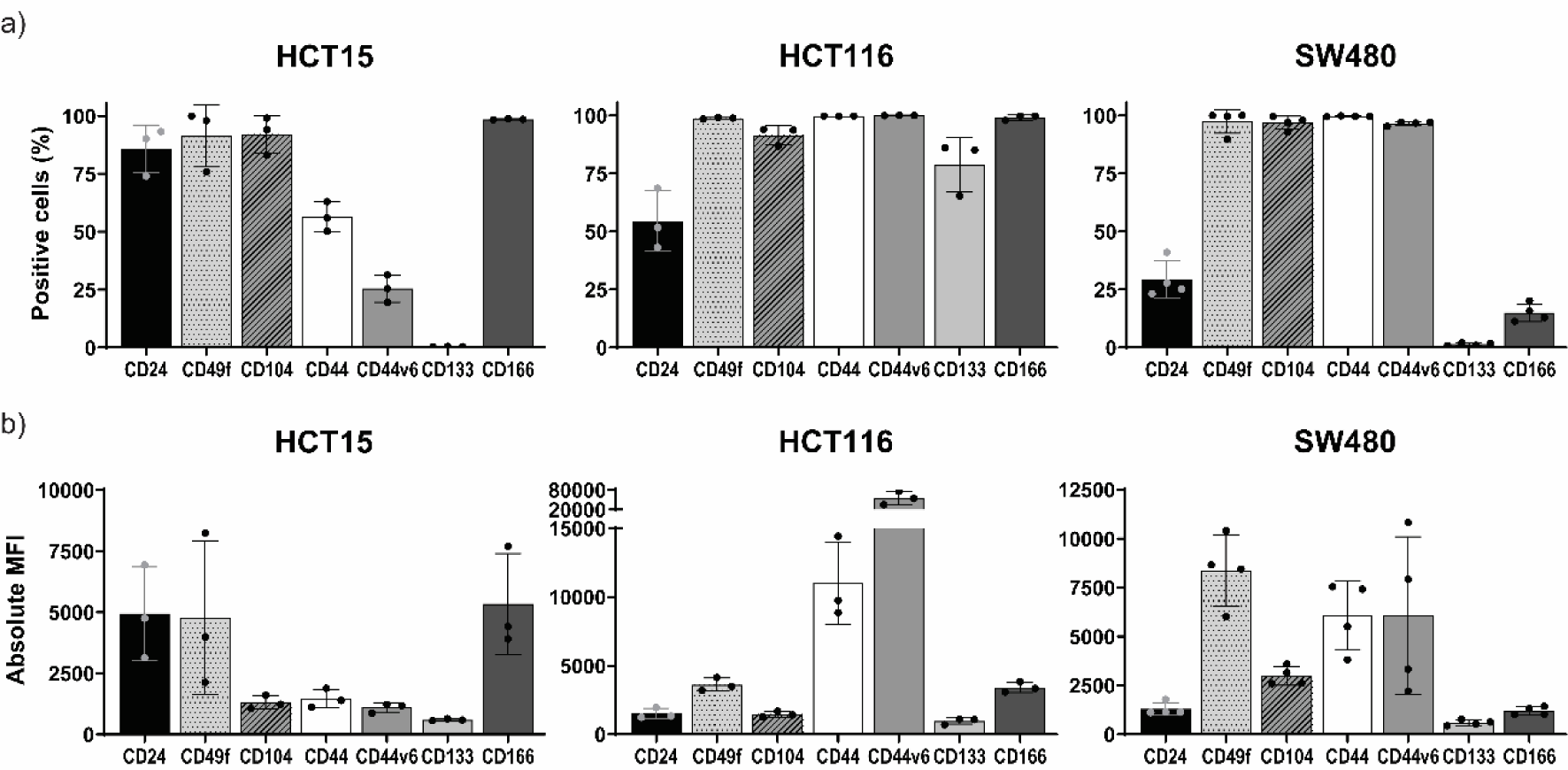
Characterization of the basal levels of stem cell marker expression in CRC cells by flow cytometry. Three different CRC cell lines with different origins and genetic profiles were selected. The most common stem cell markers were chosen to characterize the stemness potential of each cell line, by flow cytometry. Mean and standard deviation are represented in each bar. (a) Percentage of positive cells; (b) Absolute median fluorescence intensity.

### KRAS silencing leads to the upregulation of CD24 and downregulation of CD49f and CD104 across cell lines

After analyzing the basal expression of the stem cell markers for each cell line, we wanted to investigate whether KRAS was a key regulator of the expression of CRC stem cell marker signature.

First, we silenced KRAS in the three cell lines and performed flow cytometry for the markers previously described. Following KRAS silencing, there were major alterations in the expression of the stem cell markers, with the changes being more pronounced in the MFI than the percentage of positive cells (figure 2). In the HCT15 cell line, there was up-regulation of CD24 (% of positive cells and MFI) and CD44 (MFI), and down-regulation of CD49f (MFI) and CD104 (MFI). There were no significant changes observed in the expression of CD44v6, CD133, or CD166. In the HCT116 cell line, there was up-regulation of CD24 (% of positive cells), and down-regulation of CD49f (MFI), CD104 (% of positive cells and MFI), CD44 (% of positive cells and MFI), CD44v6 (% of positive cells and MFI) and CD133 (% of positive cells and MFI). There were no significant changes observed in the expression of CD166. In the SW480 cell line, there was up-regulation of CD24 (% of positive cells and MFI), CD133 (MFI), and CD166 (MFI), and down-regulation of CD49f (MFI), CD104 (MFI) and CD44 (MFI). There were no significant changes observed in the expression of CD44v6. Overall, although the cell lines suffer major alterations in the expression of the stem cell markers, only three markers displayed similar alterations in their expression pattern across the three cell lines: CD24, CD49f, and CD104. CD24 was upregulated both in the percentage of positive cells (all cell lines) and MFI (except HCT116). Inversely, CD49f and CD104 were consistently downregulated in the MFI across all cell lines.

**Figure 2:**
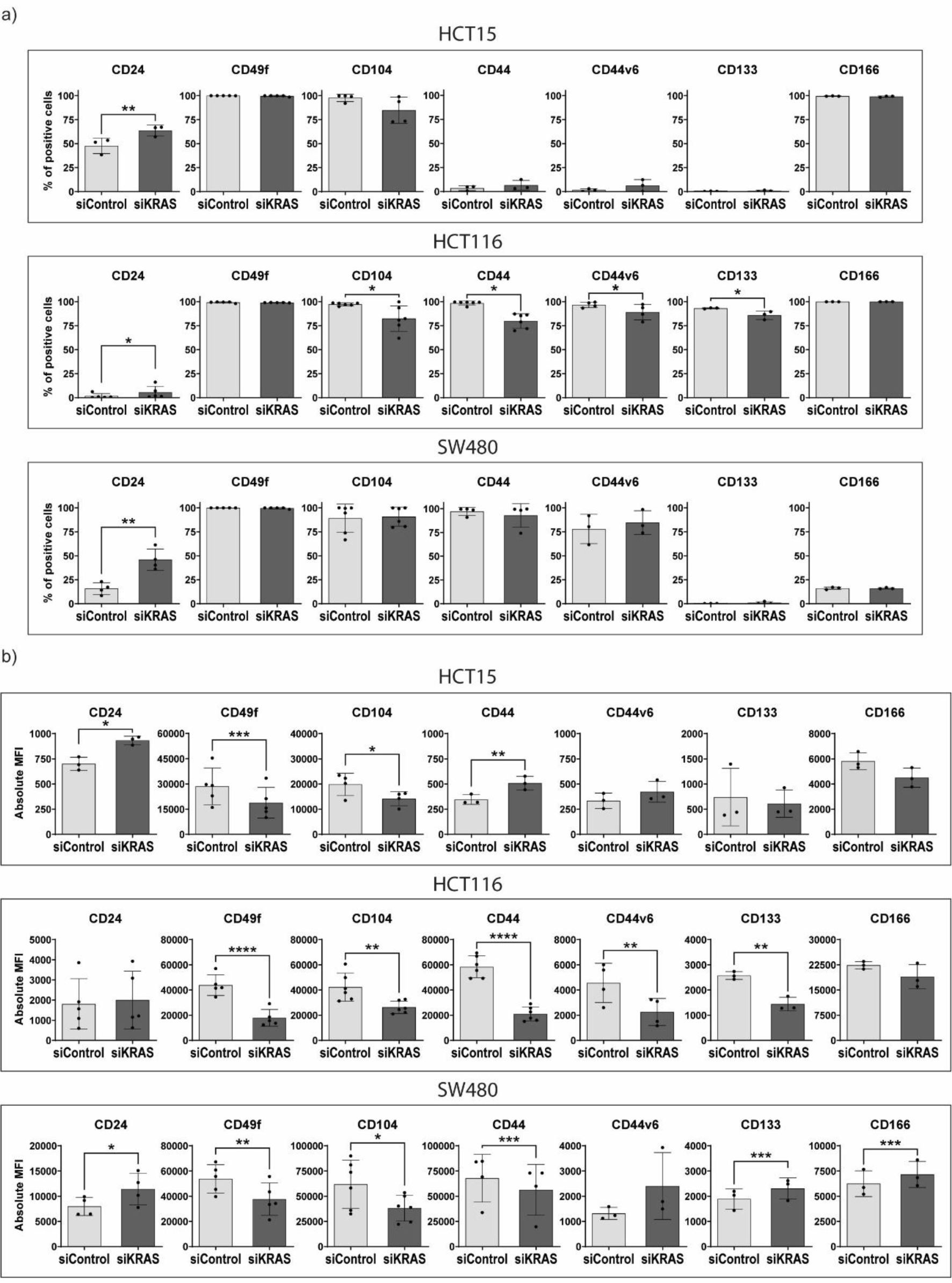
CRC stem cell marker expression upon KRAS silencing. KRAS was silenced in the three CRC cell lines (HCT15, HCT116, and SW480) and the expression of stem cell markers and integrins was analyzed by flow cytometry. For all the cell lines, normality of the data was tested using Shapiro-Wilk normality test. A one-tailed paired t-test was performed, testing for a p-value < 0,05. For the samples that did not follow normality, a Wilcoxon matched-pairs signed rank test was used. Mean and standard deviation are represented in each bar. (a) Percentage of positive cells; (b) Absolute median fluorescence intensity.

In summary, the effect of KRAS on the expression of the majority of stem cell markers is cell line dependent, except for CD24, CD49f, and CD104, whose expression across cell lines was consistently modulated upon KRAS silencing. These changes in the stem cell markers likely indicate a decrease in cancer stem cell potential upon KRAS silencing.

### Fibroblasts attenuate the capacity of KRAS silencing to regulate the expression of cancer stem cell markers

After ascertaining the regulatory role that KRAS has on the expression of stem cell markers in CRC cell lines, we sought to investigate whether fibroblasts could counteract the modulatory effects of KRAS. To accomplish this, we silenced KRAS in all the cell lines for 24 hours and subsequently treated them for 48 hours with conditioned media from activated fibroblasts (CCD-18Co normal colon cell line treated with rhTGF-β1). Strikingly, after treatment with conditioned media of activated fibroblasts, there was an attenuation in the differences previously observed between siControl and siKRAS regarding the expression of cancer stem cell markers (figure 3). For instance, in HCT15 cells, the increase in CD24, both in the % of positive cells and MFI, previously observed upon KRAS silencing, was maintained, though the reduction in CD49f and CD104 levels was lost. Additionally, CD44 levels followed an opposite trend, decreasing its expression after treatment of KRAS-silenced HCT115 cells with the fibroblast conditioned media. In the HCT116 cell line, all statistical differences between siControl and siKRAS were lost, both in the % of positive cells and the MFI. In the SW480 cell line, upon treatment with conditioned media, no statistical differences were observed between siControl and siKRAS for the markers CD24, CD49f, CD104, CD133, and CD166. However, the decrease in CD44 in KRAS-silenced cells was maintained. Contrary to what was observed in KRAS silencing alone, upon treatment with conditioned media, CD44v6 expression decreased.

**Figure 3.**
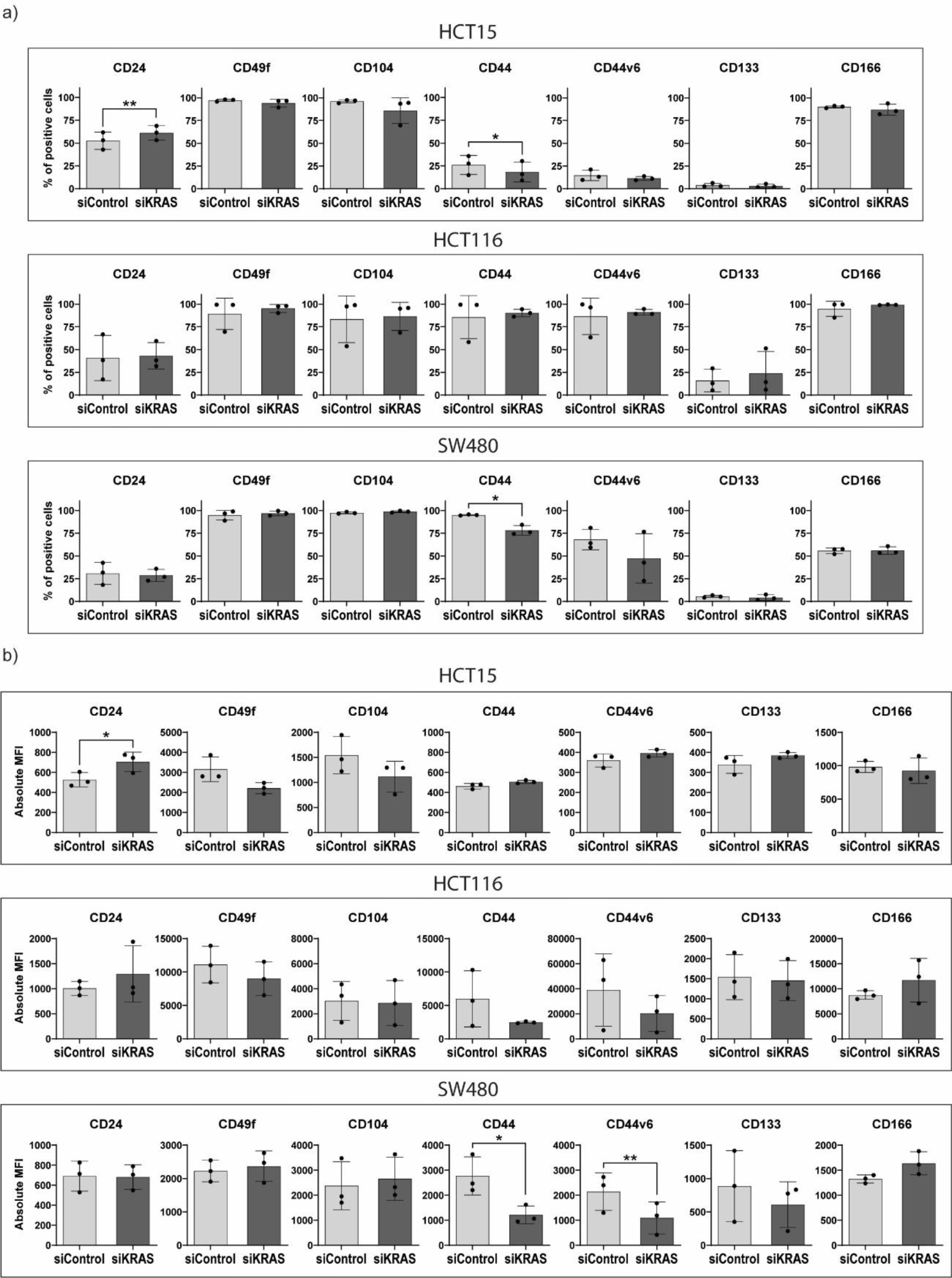
CRC stem cell marker expression upon treatment with conditioned media from fibroblasts after KRAS silencing. KRAS was silenced for 24h in the three CRC cell lines (HCT15, HCT116, and SW480). Following KRAS silencing, cells were treated for another 48 hours with conditioned media of fibroblasts activated with rhTGF-β1. Expression of stem cell markers and integrins was analyzed by flow cytometry. For all the cell lines, normality of the data was tested using Shapiro-Wilk normality test. A one-tailed paired t-test was performed, testing for a p-value < 0,05. For the samples that did not follow normality, a Wilcoxon matched-pairs signed rank test was used. Mean and standard deviation are represented in each bar. (a) Percentage of positive cells; (b) Absolute median fluorescence intensity.

Our findings highlight a significant role of fibroblast-secreted factors in influencing the response of CRC cells to KRAS silencing. In fact, they indicate that fibroblast-secreted factors are able to attenuate the effects of KRAS silencing in modulating the levels of stem cell markers in the membrane of CRC cells, unveiling another layer of intricacy in the interplay between KRAS signaling and fibroblasts.

### The sphere-forming inhibitory effect promoted by KRAS silencing is partially lost upon treatment with conditioned media from fibroblasts

To obtain a deeper understanding of how the stem cell-like phenotype is influenced by both KRAS modulation and the extrinsic signals derived from fibroblasts, we conducted an *in vitro* sphere formation assay - a technique commonly used for assessing stem cell potential based on the capacity of cells for self-renewal. For this assay, cells are plated in *anoikis* conditions. Since resistance to *anoikis* is a characteristic of stem cells, the higher the number of spheres formed, the greater the stem cell potential of that cell line.

We started by investigating the role of KRAS silencing. After silencing KRAS, CRC cancer cell lines were treated with DMEM supplemented for the sphere formation assay (B27, N2, FGF, and EGF). As depicted in figure 4a, when KRAS was silenced, the sphere-forming efficiency decreased in all cell lines, indicating a reduced stem cell-like potential across cell lines, similar to what was observed by flow cytometry. Subsequently, we tested the effect of fibroblast-secreted factors on the sphere-forming efficiency of control and KRAS-silenced cells. Surprisingly, fibroblast-conditioned media efficiently induced the formation of spheres (figure 4b), including in the KRAS-silenced conditions (except for the SW480 cell line, even though the sphere-forming efficiency of the KRAS-silenced cells in fibroblast-conditioned media is higher compared to the KRAS-silenced in control media).

**Figure 4.**
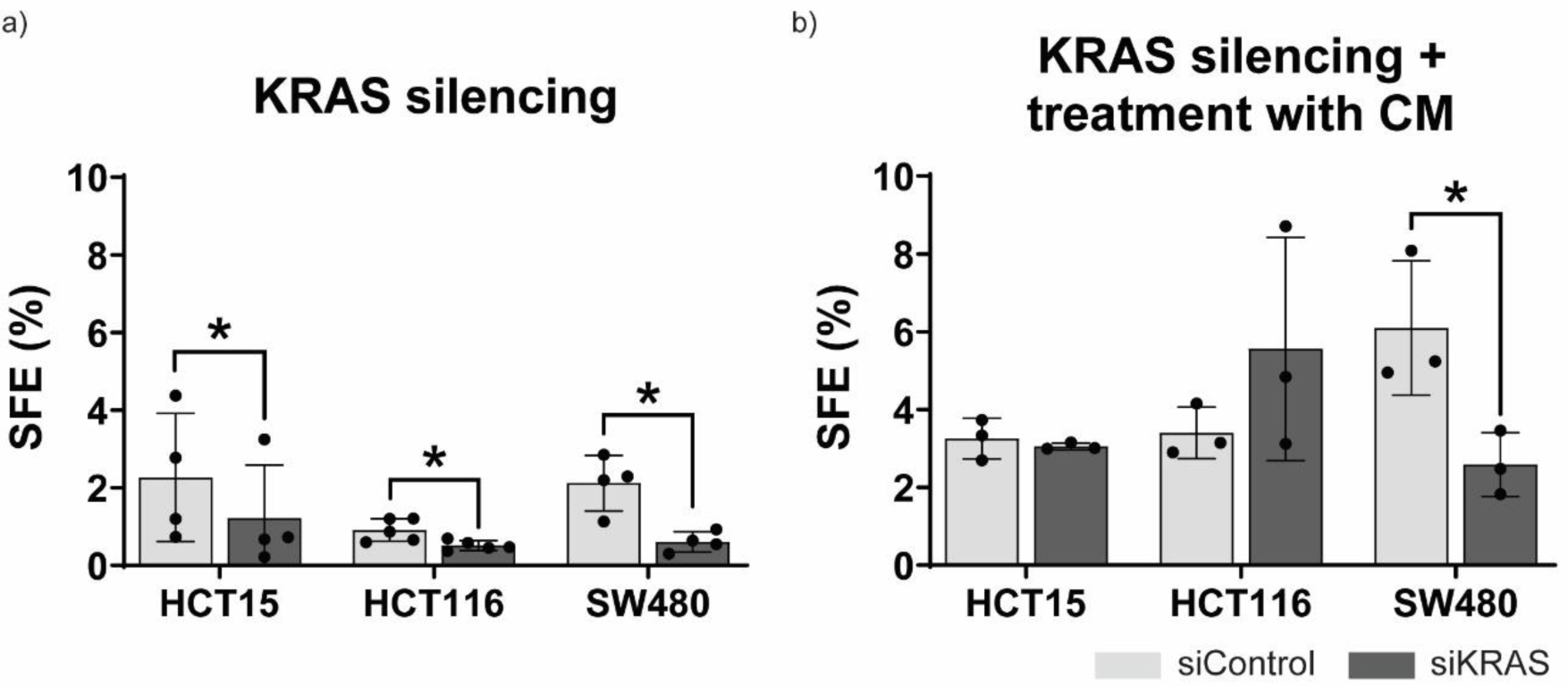
Sphere formation assay in CRC cell lines with KRAS silenced and with KRAS silenced plus treatment with conditioned media of activated fibroblasts. KRAS was silenced in the three CRC cell lines (HCT15, HCT116 and SW480). A sphere formation assay was performed using both siControl and siKRAS cells for each cell line. a) Sphere forming efficiency (SFE) percentage after treatment with DMEM supplemented for the assay (with B27, N2, FGF, and EGF; b) SFE percentage after treatment with conditioned media from activated fibroblasts (CCD-18Co cultured in DMEM with rhTGF-β1) supplemented with B27 and N2. Mean and standard deviation are shown in each bar. For all the cell lines, normality of the data was tested using a Shapiro-Wilk normality test, and a one-tailed Paired t-test was performed, testing for a p-value < 0,05.

These results suggest that treating CRC cell lines with conditioned media from fibroblasts not only enhances their self-renewal and proliferation capacity but also confers them resistance to the regulatory control imposed by KRAS silencing.

### Treatment of KRAS-silenced cells with conditioned media of activated fibroblasts upregulates proliferation, pro-tumorigenic pathways, EMT and immune system regulation

To elucidate the molecular mechanisms enabling cells treated with fibroblast-conditioned media to counteract the inhibitory effects of KRAS silencing, we performed RNASeq analysis. We decided to focus on the HCT116 cell line because it showed the biggest sphere-forming efficiency increase when comparing siKRAS and siControl upon treatment with conditioned media from activated fibroblasts. By comparing KRAS-silenced spheres treated either with conditioned media of activated fibroblast (supplemented with B27 and N2) or with control media (DMEM supplemented with B27, N2, FGF, and EGF), we aimed to identify the specific genes and pathways influenced by fibroblast-derived factors, thereby revealing crucial players in this complex cellular interplay. We analyzed our data set using Gene Set Enrichment Analysis (GSEA) computational method and its Molecular Signatures Databases (MSigDB) to look at gene set collections. KRAS-silenced cells treated with conditioned media exhibit a distinctive gene expression profile in comparison to cells treated with control media (Figure 5a and supplementary table 1). Then, we analyzed the hallmarks of molecular signature that is represented in Figures 5b and c. For cells treated with conditioned media of activated fibroblasts, the top up-regulated pathways are mainly associated with cell cycle control (E2F targets, G2-M checkpoints, MYC targets, mitotic spindle pathways), EMT, immune system regulation (IL6, IL2, interferon-gamma, complement pathways) and several other signaling pathways (KRAS, NOTCH, TGF-β and WNT) that can be associated with carcinogenesis. On the other hand, the top up-regulated pathways in cells treated with control media were mainly associated with metabolism (cholesterol, fatty acids, and xenobiotic metabolism), cell stress (apoptosis and hypoxia, TNF-α and mTOR signaling pathways), and DNA damage response (unfolded protein response and P53 pathways).

**Figure 5.**
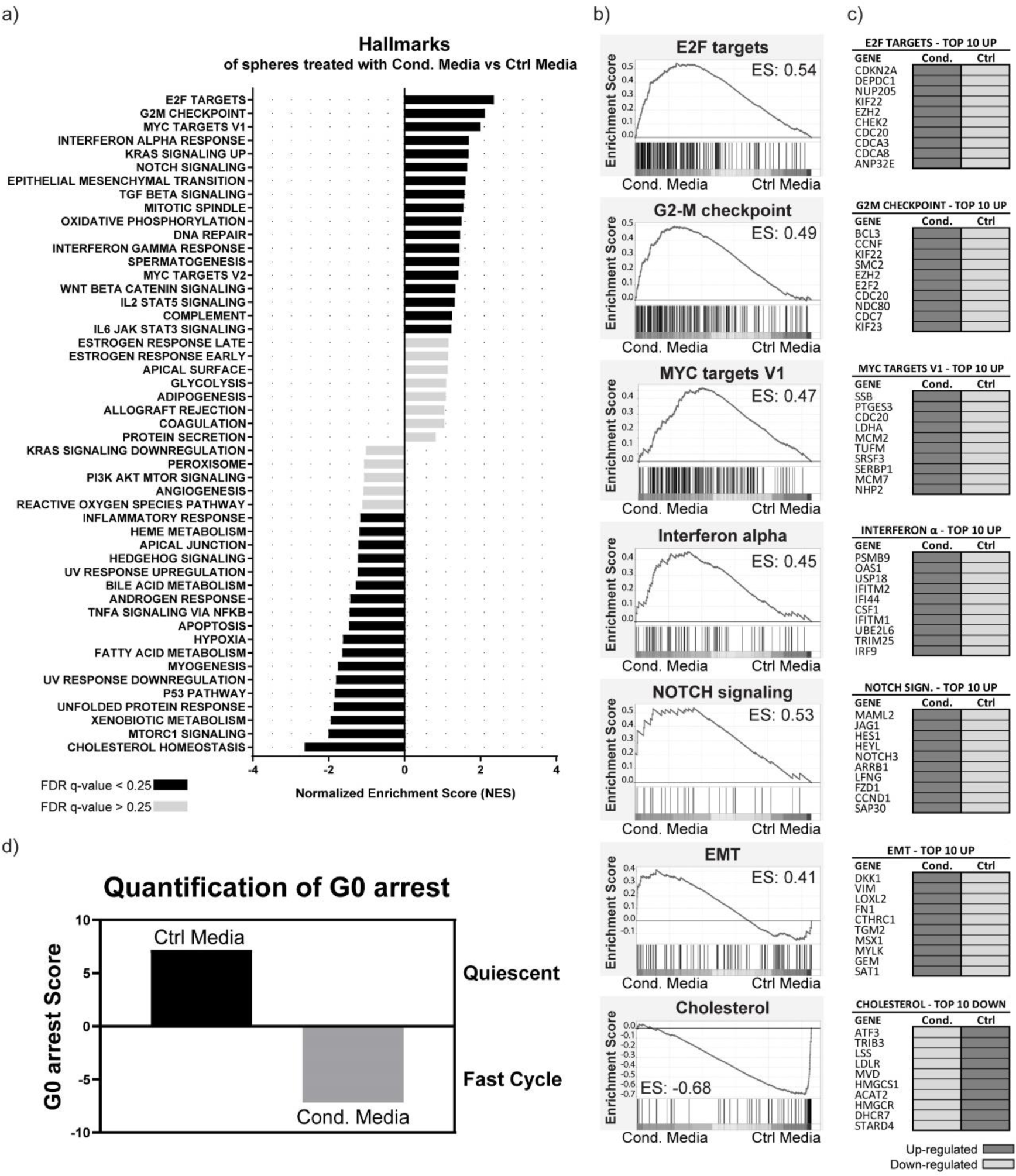
RNAseq analysis performed in GSEA and evaluation of G0 arrest transcriptional signature profiling. RNA-Seq was performed on a pool of three experiments of KRAS silenced HCT116 spheres treated with conditioned media of activated fibroblasts supplemented with B27 and N2 (Cond. Media) or control media - DMEM without phenol red supplemented with FGF, EGF, B27, and N2 (Ctrl Media). a) Results of GSEA Hallmark analysis showing enriched gene sets. Black bars indicate significant enrichment at false discovery rate (FDR) < 25% while grey bars represent gene sets with FDR > 25%. A positive normalized enrichment score (NES) value indicates enrichment in the cells treated with conditioned media of fibroblasts whereas a negative NES indicates enrichment in the cells treated with control media. b) Enrichment plots for the most relevant data sets enriched in GSEA Hallmark analysis, showing the profile of the running enrichment score (ES) and positions of gene set members on the rank-ordered list. c) Tables showing the top 10 enriched genes in each data set. Dark grey indicates up-regulation of the gene while light grey color indicates down-regulation. (d) Quantification of G0 arrest by the method described by (Wiecek et al., 2023). Positive values indicate that cells are in a quiescent state, whereas negative values signify cells are in a proliferative cycle.

Given the upregulation of numerous genes associated with cell cycle control in cells treated with conditioned media from fibroblasts, we aimed to determine whether these cells were indeed more proliferative than the cells treated with control media. Wiecek et al., 2023 developed a methodology to determine if cells are in a state of G0 arrest that encompasses quiescence, senescence, and dormancy or if, alternatively, they are in a rapid cell cycle progression state similar to that associated with stem cells and their self-renewal capacity. Based on the approach described in their research, we analyzed the G0 arrest transcriptional signature of our cells. As represented in figure 5d, the analysis of the G0 arrest transcriptional signature revealed that KRAS silenced cells treated with conditioned media of fibroblasts were in a state of fast cell cycling (negative G0 score) while the KRAS silenced cells in control media were in a quiescent state (positive G0 score).

These results suggest that fibroblast-secret factors overcome the growth inhibitory effect of KRAS silencing, increasing their proliferative potential and transitioning to a state closer to a mesenchymal phenotype through the upregulation of EMT and stemness-related molecules.

## Discussion

In a time of intense search for therapeutic strategies to efficiently target KRAS mutant tumors, our *in vitro* study uncovers a pivotal mechanism to bypass KRAS silencing, independent of KRAS and orchestrated by cancer-associated fibroblast-derived factors that trigger cancer stem cell activity, proliferation, EMT, modulation of immune response and up-regulation of tumorigenic pathways.

Located downstream of many cell surface receptors, KRAS is considered a central regulator of intracellular signaling in response to extracellular stimuli (Dias Carvalho, Martins, et al., 2022; Dias Carvalho, Mendonça, et al., 2022; Huang et al., 2021). Therefore, it is associated with the regulation of diverse cancer cell activities, including the induction of a stem-like phenotype (Chippalkatti & Abankwa, 2021). Targeting CSCs to avoid cancer recurrence is one of the most significant challenges in oncology (Prieto-Vila et al., 2017). Understanding the factors and circumstances that regulate CSCs and that govern the development of recurrence is crucial to improve treatment efficacy (Prieto-Vila et al., 2017).

In our study, we looked at stem cell markers by flow cytometry in CRC cell lines harboring KRAS mutations. Our results show that, although stem cell markers are altered in a cell line-dependent manner, KRAS silencing was consistently associated with an up-regulation of CD24 and down-regulation of CD49f and CD104 in all the cell lines studied. Although CD24 upregulation has been previously correlated with tumor progression, invasiveness, differentiation, and chemotherapy resistance (D. Choi et al., 2009; Paschall et al., 2016; Yeo et al., 2016) some other studies hint that loss of CD24 expression might be more associated with poorer outcomes (Ahmed et al., 2009) while others suggest that CD24 is a good prognosis marker in CRC, being downregulated in stage IV colorectal adenocarcinoma (Yeo et al., 2016). Therefore, the role of CD24 on CRC and CRC stem cell phenotype is still not consensual.

The complex CD49f/CD104 is important for cell-matrix interaction and is usually found over-expressed in CRC. Although the mechanisms that control its expression are still not fully known, transcription factors, such as MYC, have been implicated in this process. Coincidentally, CD49f/CD104 can increase cell proliferation through the activation of the Wnt/β-catenin pathway, a regulator of stem cell homeostasis in the intestinal crypts and an upstream effector of MYC (Beaulieu, 2020). It was demonstrated that depletion of integrin CD49f/CD104 reduces the invasive and migratory capabilities of the cancer cells (S. H. Choi et al., 2022). Our findings concur with existing literature and suggest a phenotypic shift in CRC cells upon KRAS silencing towards a less stem cell-like and less aggressive state.

However, when we treat these KRAS-silenced cells with conditioned media from fibroblasts, we see an alteration in the KRAS-modulated stem cell marker expression profile. Our findings align with the observations made by Colak & Medema, 2016 who stated that CAF-secreted factors can lead to the de-differentiation of non-CSCs. These alterations in the membrane stem cell markers might not be, however, direct indicators of real stemness potential since Vermeulen et al., 2008 showed that CSC activity is not directly associated with the expression of CSC biomarkers. Therefore, we decided to perform a sphere formation assay, which allowed us to confirm that, indeed, fibroblasts, even in KRAS-silenced cells, can potentiate the stem cell potential of CRC cells. It is important to note that the sphere formation assay, although a useful tool that provides clues about stemness potential in cell lines serves only as an *in vitro* model (Pastrana et al., 2011). Further *in vivo* studies analyzing the response of CRC cells to KRAS-targeted inhibitors within a fibroblast-rich or poor microenvironment will confirm whether the observed results genuinely represent stem cell properties.

Our RNAseq results further reinforced the role of fibroblasts as mediators of resistance to KRAS silencing, detailing the upregulation of cell cycle control, EMT, immune system regulation, and cancer-associated pathways such as KRAS, NOTCH, TGF-β, MYC, and WNT as the main pathways induced by fibroblast-derived factors. Specifically, E2F, MYC, and G2M pathways, frequently found to be downregulated upon KRAS inhibition (Hallin et al., 2020; Tammaccaro et al., 2023) ranked on the top three upregulated biological processes found in KRAS-silenced cells stimulated with fibroblast-conditioned media.

Stem cell-like phenotype and EMT are deeply intertwined, as cells undergoing EMT can acquire CSC properties, while CSCs themselves can undergo EMT to facilitate metastasis (Fiori et al., 2019). It has been shown that conditioned media of CAFs is enriched in TGF-β1 and, through the activation of the canonical TGF-β pathway, can induce EMT (Zhuang et al., 2015). Furthermore, TGF-β signaling has been correlated with CRC subtypes with worse prognosis and increased relapse. Conditioned media of CAFs is also enriched in other factors such as fibroblast growth factor (FGF), interleukin-6 (IL-6), hepatocyte growth factor (HGF), osteopontin (OPN), and stromal-derived factor-1α (SDF1) (Fiori et al., 2019). The latter three have been shown to have an important role in modulating cancer cells into a more stem cell-like phenotype, particularly through the Wnt/β-catenin signaling pathway, which can also increase CD44v6 expression. IL-6, on the other hand, plays a pivotal role in immune system regulation by promoting a chronic inflammatory environment. Interestingly, IL-6 can also control the activation of NOTCH, a particularly relevant pathway because it is responsible for the self-renewal of intestinal stem cells, CSC maintenance, TGF-induced EMT, and resistance to therapy (Deshmukh et al., 2021; Ebrahimi et al., 2023; Fiori et al., 2019). In our results, one of the top up-regulated genes in the NOTCH signaling pathway is NOTCH3, whose aberrant expression is a recurring finding in human cancer tissues. In CRC, NOTCH3 expression increases with tumor staging and is correlated with worse prognosis, poor overall survival, and CMS4 tumors, which are CAF-enriched (Varga et al., 2020; Xiu et al., 2021). NOTCH3 also plays a role in supporting cancer stemness and resistance to therapy (Xiu et al., 2021), which can be circumvented by the use of NOTCH inhibitors since they reduce the expression of stem cell markers and improve response to chemotherapy and radiotherapy (Ebrahimi et al., 2023). Therefore, we suggest that NOTCH inhibitors, such as gamma-secretase, could be tested in combination with KRAS-targeting therapies to try to overcome the maintenance of stemness produced by CAFs and improve the current problem of resistance to the therapy.

Overall, our work shows that KRAS-silenced CRC cells lose stem cell-like properties and undergo a quiescent-like state. Our results award fibroblasts a key role in overcoming the antitumorigenic effects of KRAS silencing, implicating them as potential mediators of resistance to KRAS-targeted therapy. On the one hand, fibroblast-rich CRC, such as those belonging to the CMS4, may display cancer cell-extrinsic innate resistance to KRAS inhibition, highlighting fibroblast infiltration as a potential biomarker of lack of response to KRAS inhibition. In this case, fibroblasts aid cancer cells with pro-tumorigenic signals that allow them to circumvent the absence of KRAS oncogenic signaling, reinstalling growth and de-differentiation. On the other hand, our results, together with our previous observations showing that KRAS-silenced HCT116 cells promote fibroblast migration while retaining the capacity to activate them (Dias Carvalho, Mendonça et al., 2022), hint at a potential mechanism of acquired resistance positive driven by a positive feedback loop: KRAS-inhibited fibroblast-poor CRC recruit fibroblasts and activate them into CAFs which in turn sustain CRC growth in a KRAS-independent manner.

In conclusion, our findings underscore the necessity to rethink the current therapeutic strategies to target KRAS mutant CRCs. Targeting KRAS alone appears to be insufficient, and CRC patients could benefit from combined therapies where not only we target KRAS but also the CAFs present in the tumor microenvironment. Our study already highlights some potential CAF-derived molecular targets worthy of being explored therapeutically.

## Author Contributions

Conceptualization: S.M.O., S.V. and M.J.O.; Formal analysis: S.M.O., P.O. and A.R.; Funding acquisition: S.V.; Investigation, S.M.O.; Methodology: S.M.O., P.D.C., A.R., P.O., A.R., F.M., A.L.M., J.C. and A.R.; Project administration: S.V.; Resources: S.V.; Supervision: M.J.O. and S.V.; Visualization: S.M.O.; Writing-original draft: S.M.O.; Writing-review and editing: S.M.O. and S.V. All authors have read and agreed to the published version of the manuscript.

## Acknowledgments

The authors acknowledge the support of the i3S Genomics Scientific Platform, in particular Ana Mafalda Rocha. The authors also acknowledge André Maia from the BioSciences Screening i3S Scientific Platform, and Catarina Meireles and Emilia Cardoso from the Translational Cytometry unit (TraCy) i3S Scientific Platform.

## Abbreviations

BSA: Bovine serum albumin
CAFs: Cancer associated fibroblasts
CRC: Colorectal cancer
CSCs: Cancer stem cells
ECL: Enhanced chemiluminescence
EDTA: Ethylenediaminetetraacetic acid
EGF: Epidermal growth factor
EMT: Epithelial-mesenchymal transition
ES: Enrichment score
FBS: Fetal bovine serum
FDS: False discovery rate
FGF: Fibroblast growth factor
GSEA: Gene set enrichment analysis
HRP: Horseradish peroxidase
KRAS: Kirsten rat sarcoma virus
MFI: Median fluorescence intensity
NES: Normalized enrichment score
P/S: Penicillin–streptomycin
PBS: Phosphate-buffered saline
rhTGF-β1: Recombinant human transforming growth factor beta 1
RIPA: Radioimmunoprecipitation assay
SFE: Sphere forming efficiency
siControl: Control cells
siKRAS: KRAS silenced cells
siRNA: Small interfering RNAs

## Supplementary Information

Supplementary Table 1-Differentially expressed genes with 2.5x fold change in the RNAseq analysis.

Supplementary Figure 1-WB showing activation of fibroblasts, KRAS silencing in the different experiments, and gating strategy for flow cytometry.

