## Supplementary Figure 1 for "Unveiling the Mechanisms to Bypass KRAS Inhibition: *In Vitro* Insights into the Influence of Fibroblast-Secretome"

### Figure S1

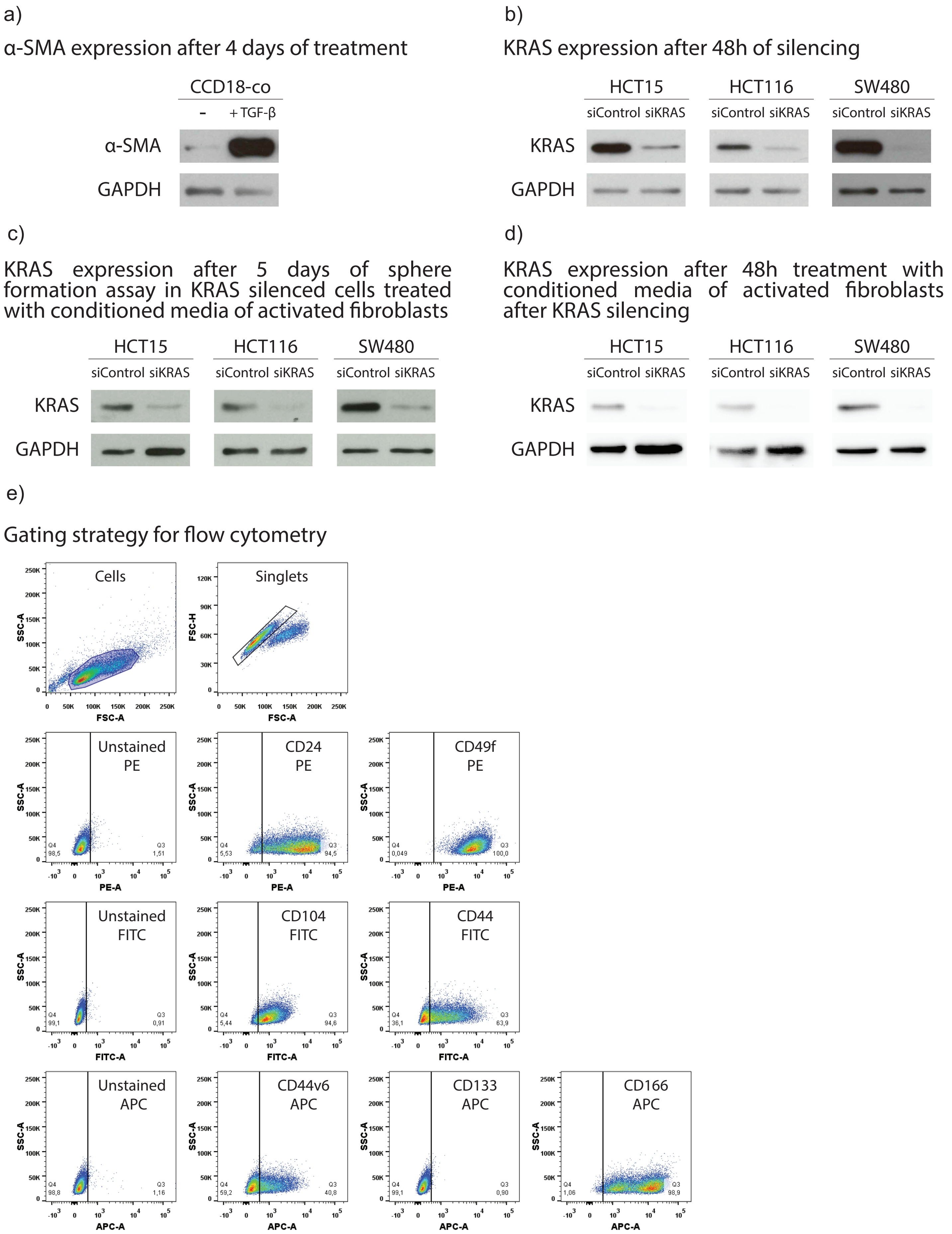


***Figure S1: Validation of the activation of fibroblasts and KRAS silencing by Western Blot.*** *GAPDH was used as a loading control for all the experiments. (a) Protein of normal colon fibroblast cell line CCD-18co was collected after 4 days of culture in DMEM either without (-) or with addition of TGF-β (+TGF-β) which activates the fibroblasts. α-SMA is used to determine the activation state of fibroblasts since activated fibroblasts show an increased expression of this protein. (b) Protein was colected after 48h of KRAS silencing in the three different colorectal cancer cell lines. (c) Protein was colected after after 5 days of sphere formation assay in KRAS silenced cells treated with conditioned media from fibroblasts. (d) Protein was colected after 48h of KRAS silencing plus treatment with conditioned media of fibroblasts. e) Example of the gating strategy used to identify cell populations in FlowJo software (HCT15 cell line is shown in the example).*
